# New Host and Geographic Range Expansion of *Lemuricola nycticebi* in the Bengal Slow Loris

**DOI:** 10.64898/2026.01.09.698086

**Authors:** Joytu Kumar Mondal, Md. Rajiur Rahaman Rabbi, Mahir Anjum, M Farhan Sadid Chowdhury, Ruhel Ahmed, Atiya Ibnat Lamia, Meherab Hossain, Atika Angum Miti, Chonchol Guala, Ibrahim Amin, Fahimuzzaman Nobel, Shahriar Caesar Rahman, Christian Roos, Tanvir Ahmed

## Abstract

**Purpose:** The Bengal slow loris (*Nycticebus bengalensis*) is a globally Endangered nocturnal primate with limited health data across its South and Southeast Asian range.

**Methods:** During necropsies of two deceased individuals from northeastern Bangladesh, numerous live adult pinworms were recovered.

**Results:** Morphological examination combined with mitochondrial *cox1* gene sequencing identified the parasites as *Lemuricola (Protenterobius) nycticebi*. Both hosts exhibited severe gastrointestinal and peritoneal lesions consistent with high-intensity helminth infection. This represents the first confirmed record of *L. nycticebi* in the Bengal slow loris and extends the parasite’s known geographic range by approximately 3,400 km.

**Conclusion:** Our findings highlight an overlooked parasitic threat to slow lorises. Given that many Bengal slow lorises are rescued or confiscated from the illegal wildlife trade and forest-edge areas in Bangladesh every year and subsequently released into the wild, the results underscore the need for systematic health assessments, including parasitic screening, before repatriation into the wild.

## 1. INTRODUCTION

Parasites represent some of the most diverse and ecologically influential organisms on the planet, shaping host population dynamics, altering community structure, and driving evolutionary processes [1,2,3]. Globally, parasitic infections have substantial impacts on human and animal health, influencing survival, reproduction, and overall ecosystem stability [2,3]. As biodiversity declines and ecosystems continue to undergo anthropogenic change, understanding parasite diversity and distribution has become increasingly important for both conservation biology and disease ecology [1,2,4,5].

Wildlife parasites, in particular, play complex and often overlooked roles in natural systems [2,6. While many parasites coexist benignly with their hosts, others can contribute to morbidity and mortality, especially when environmental stressors, habitat fragmentation, or human–wildlife interactions disrupt long-standing host–parasite relationships [2,5,7,8,9]. Detailed knowledge of parasite fauna in wildlife species is therefore crucial not only for disease surveillance but also for informed management of threatened taxa, rehabilitation programs, and translocation initiatives [2,9,10]. Despite this importance, parasite diversity in many wildlife groups remains poorly characterized, especially in regions with limited research infrastructure [3,12].

Primates host an extensive array of parasitic organisms, including protozoans, helminths, and arthropods, reflecting their ecological diversity, complex social systems, and broad biogeographic distribution [8,13,14]. Parasitological studies in primates have provided crucial insights into host behaviour, nutrition, evolutionary history, and conservation issues [15,16,17]. However, research efforts have been uneven, with substantial attention given to African great apes and macaques, while many Asian primates, particularly nocturnal and elusive species, have received far less investigation [8,18,19]. This knowledge gap limits our ability to assess health risks, manage rehabilitation and reintroduction programs, conservation breeding centers and understand parasite transmission at the human–wildlife interface [8,17,19,20,21].

Slow lorises (*Loris* spp., *Nycticebus* spp., *Xanthonycticebus* spp.), the only nocturnal strepsirrhines in Asia, exemplify this research gap [22]. Distributed across South and Southeast Asia, slow lorises face severe threats from habitat loss, hunting, and the illegal wildlife trade, making them among the most frequently confiscated primate species in the region [23,24]. Despite their high conservation priority, information on the parasite fauna of slow lorises remains scarce or entirely lacking [22,25]. For example, the Bengal slow loris (*Nycticebus bengalensis*), classified as Endangered and subject to intense trafficking across South and Southeast Asian countries [27,28,29], remains entirely unstudied with respect to its parasitic diversity. Particularly in Bangladesh, every year, many primates, especially Bengal slow lorises, are rescued from forest-fringe areas and personal collections or confiscated from the illegal wildlife trade [27,28,29,30]. Often, such animals are subsequently released to the wild without considering the risks of parasitic and disease transmission, their survival potential, and other negative impacts on both translocated individuals and recipient ecosystems [29,31].

Among the limited helminths recorded in slow lorises, pinworms of the genus *Lemuricola* (family Oxyuridae) are particularly noteworthy, as they are specific to strepsirrhines [22,25]. Pinworms are cryptic nematodes exceptional for their ability to infect a wide range of hosts. Infections are often asymptomatic but can cause inappetence, progressive weight loss, and lethargy. One species, *Lemuricola nycticebi*, was first described from the Philippine slow loris (*Nycticebus menagensis*) in Sarawak, Malaysian Borneo [32], and later redescribed from a Sunda slow loris (*Nycticebus coucang*) in Peninsular Malaysia [33]. Some insights into its evolutionary history were also reported from free-living Philippine slow lorises in Sabah, Malaysian Borneo [25], but its host range and geographic distribution remain poorly understood. To address this knowledge gap, we document *L. nycticebi* infections in the Bengal slow lorises for the first time, supported by both morphological assessment and genetic confirmation of specimens obtained in Bangladesh.

## 2. METHODS

### 2.1 Gross-pathological examination and sample collection

This study was conducted at the Jankichora Wildlife Rescue Center, located within Lawachara National Park in northeast Bangladesh. The center is operating wildlife rehabilitation and specialized wildlife clinic facilities and is jointly managed by the Bangladesh Forest Department and the non-governmental organization Creative Conservation Alliance. Every year, numerous rescued or confiscated wild animals, often suffering from severe injuries due to electrocutions and road accidents, are brought to the center by the forest officials.

For this study, worm specimens were collected from two deceased Bengal slow lorises brought to the clinic by the regional office of the Bangladesh Forest Department. One individual died following a collision with vehicles on the highway passing through Lawachara forest, and the interval was approximately 3 hrs. The second individual presented with a clinical history of bloody-mucous diarrhoea; however, its origin was unclear, and it was suspected to be either a confiscated animal from illegal wildlife trafficking or a locally rescued individual from outside the forest area. We conducted a thorough gross examination of both loris carcasses immediately after arrival. The examination includes external inspection for signs of trauma, injury, or abnormal lesions. Internal organs were examined following standard necropsy procedures to detect any pathological changes such as haemorrhage, congestion, necrosis or parasitic lesions [34]. The gastrointestinal tract was opened systematically from the esophagus to the rectum to assess the presence of worms, inflammation, or other abnormalities [35]. Live worms were carefully collected from the opened intestinal tract and immediately transferred into containers containing normal saline. Notable findings were photographed and documented subsequently. Finally, the collected specimens were transferred to the Interdisciplinary Institute for Food Security (IIFS) of Bangladesh Agricultural University for detailed phenotypic examination and genetic profiling.

### 2.2 Morphological identification

For microphotography, worms were initially cleared in a glycerol-ethanol solution first and subsequently mounted on glass slides with a 50% glycerol aqueous solution and studied at different magnifications with an Olympus (Olympus Corporation, Tokyo, Japan) light microscope BX53 equipped with an Olympus DP27 microscope digital camera and cellSens imaging software (Olympus Corporation, Tokyo, Japan). Before genetic confirmation, we initially identified the worms up to species level based on their standard and comparable morphological characters described in the previous studies [22,25,26,32,33].

### 2.3 Genetic profiling

The genomic DNA was extracted from (n = 12) adult worms using the TIANamp Genomic DNA Kit (TIANGEN Biotech (Beijing Co., Ltd.)) according to the manufacturer’s instructions. Subsequently, the extracted genomic DNA was subjected to PCR amplification targeting the 845bp-long mitochondrial cytochrome c oxidase subunit 1 (*cox1*) gene using primers StrCoxAfrF 5’-GTGGTTTTGGTAATTGAATGGTT-3’ and X pr-b 5’-AGAAAGAACGTAATGAAAATGAGCAAC-3’. PCR reactions were performed in a total volume of 25 μl reaction volume containing 12.5 μl Premix Taq™ (TaKaRa Bio, Japan), 1.5 μl (10 µM) of each primer 4.5 μl molecular grade water, and 5 μl genomic DNA. The cycling profile for the *cox1* region consisted of an initial denaturation at 94 °C for 2 min, followed by 20 cycles at 94 °C for 60 s, 55 °C for 60 s, 68 °C for 60 s and a final extension at 68 °C for 7 min (Hasegawa et al., 2012). The PCR products were run on an agarose gel (1.5%) stained with Midori Green Advance (NIPPON Genetics EUROPE), excised from the gel and then purified with the FavorPrep™ Gel/PCR Purification Mini Kit (Biotech Corp., USA). Purified PCR products were sent to Genecreate Biotech, China, for Sanger [37] sequencing using both amplification primers.

All generated sequences were annotated and trimmed, and then subjected to BLASTn search. Phylogenetic analysis was conducted together with orthologous sequences of closely related species identified by BLAST. A multiple sequence alignment was performed using MUSCLE in MEGA version 11 [38], followed by manual trimming to remove poorly aligned regions. Phylogenetic modelling and tree visualization were carried out in MEGA 11 using the Maximum Likelihood (ML) method [38]. The robustness of the phylogenetic tree was validated by running the analysis on 1000 bootstrapped input datasets and cross-referencing it against the Tamura-Nei substitution model.

## 3. RESULTS

### 3.1 Parasito-pathological changes

The post-mortem examination of the two deceased slow lorises (Fig. 1A-B) revealed significant parasitic and pathological alterations. Numerous live worms were observed within the inner peritoneum (Fig. 1C) and throughout the large intestinal contents (Fig. 1D), as well as embedded in the peritoneal–diaphragmatic wall (Fig. 1E). The intestinal organs exhibited extensive hemorrhages (Fig. 1F), and multiple hemorrhagic patches were evident on the mucosal surface of the large intestine (Fig. 1G).

**Fig. 1.**
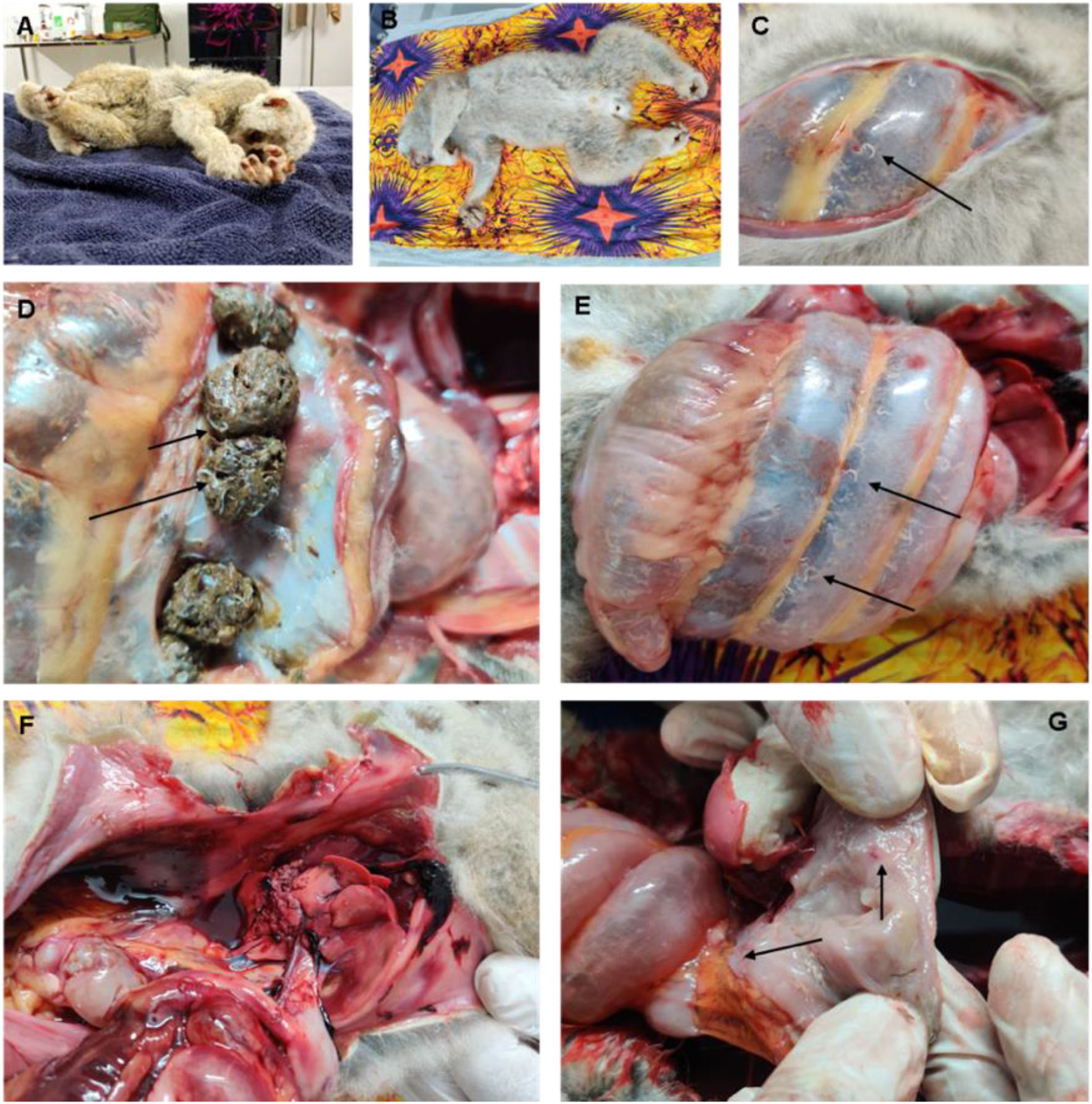
Necropsy of two deceased Bengal slow lorises at the wildlife clinic in Jankichora Wildlife Rescue Center, Northeast Bangladesh. (A-B) Dead carcasses of the animals sent for necropsy. (C) Live worms in the inner peritoneum. (D) Live worms in the large intestinal contents. (E) Pinworms in the peritoneal-diaphragmatic wall of the slow loris. (F) Extensive hemorrhages in the intestinal organs. (G) Hemorrhagic patches in the large intestinal mucosa.

### 3.2 Micromorphology of the worms

The body of the specimen is slender, filiform, and whitish, tapering gradually towards the posterior end (Fig. 2A). The anterior region bears three prominent buccal lips surrounding the mouth, leading into a muscular pharynx (Fig. 2B). The esophageal region comprises a well-defined corpus, a narrow isthmus, and a prominent posterior bulb (Fig. 2C–D). A lateral ala is visible along the anterior portion of the body. The gravid uterus contains numerous ovals, rice-seed-shaped eggs with smooth shells (Fig. 2E). The cuticle is finely striated (Fig. 2F) and the posterior part of the body houses a distinct rectum (Fig. 2G). The tail is long, slender, and filiform, extending posteriorly (Fig. 2H).

**Fig. 2.**
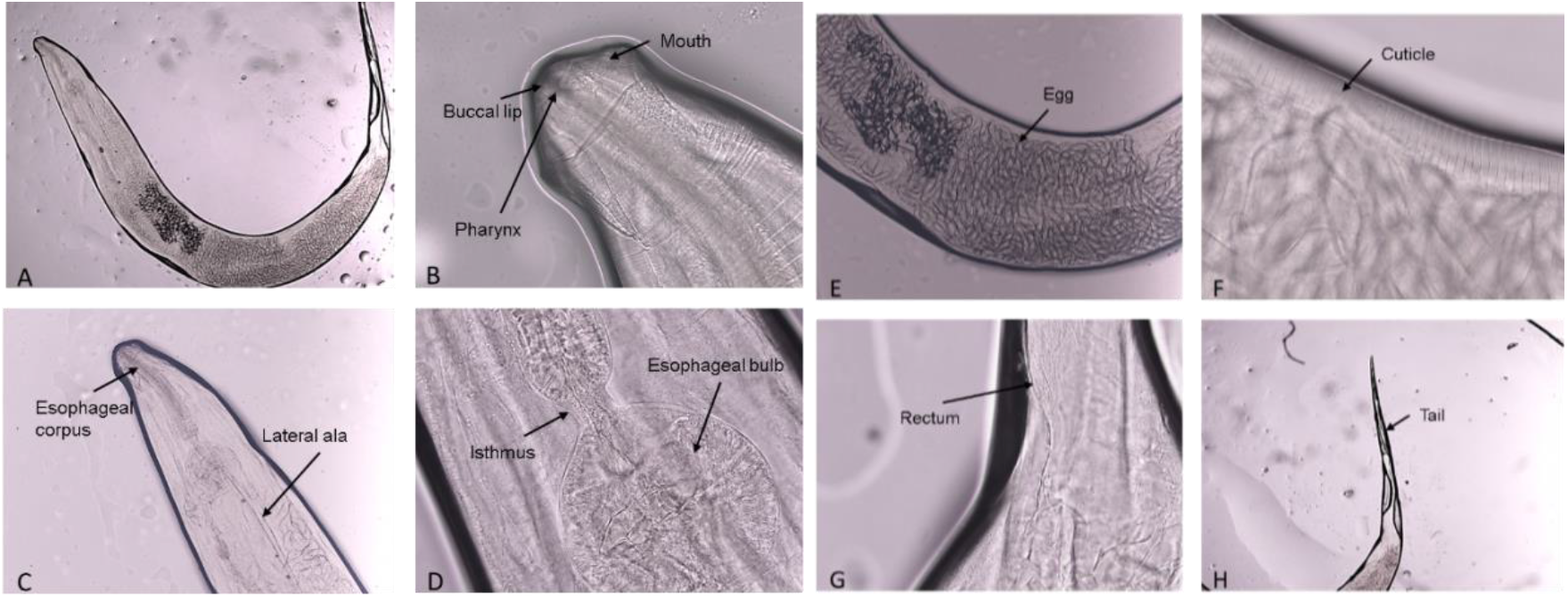
Micromorphology of adult *L. nycticebi* (Female). (A) Adult (female) worm (10X). (B) Three buccal lips and pharynx in the mouth region (40X). (C) Esophageal corpus and lateral ala in the anterior part of the adult worm (40X). (D) Isthmus and a prominent bulb in the esophageal region (40X). (E) Oval (rice seed) shaped egg in gravid worm (40X). (F) Cuticle of adult worm (40X). (G) Rectum in the posterior part of the worm (40X). (H) Filiform tail of adult (female) worm (40X).

Based on the abovementioned morphological features observed under the light microscope and key characters typical of strepsirrhine-associated oxyurids (nematode pinworms), the specimens were initially identified as adult *L. nycticebi*.

### 3.3 Molecular identification

The mitochondrial DNA of the worms obtained from both slow lorises was successfully amplified (845bp) and sequenced (Figure S1). The two sequences generated from the deceased lorises (GenBank accession numbers: PX506060 and PX506061) showed 99.88% and 99.76% nucleotide identity, respectively, with the Malaysian slow loris pinworm (*L. nycticebi*; LC416076.1 and LC416077.1). Phylogenetic analysis revealed that both sequences clustered together in a well-supported monophyletic clade with the Malaysian *L. nycticebi* and were positioned within the *Lemuricola* group (Fig. 3).

**Fig. 3.**
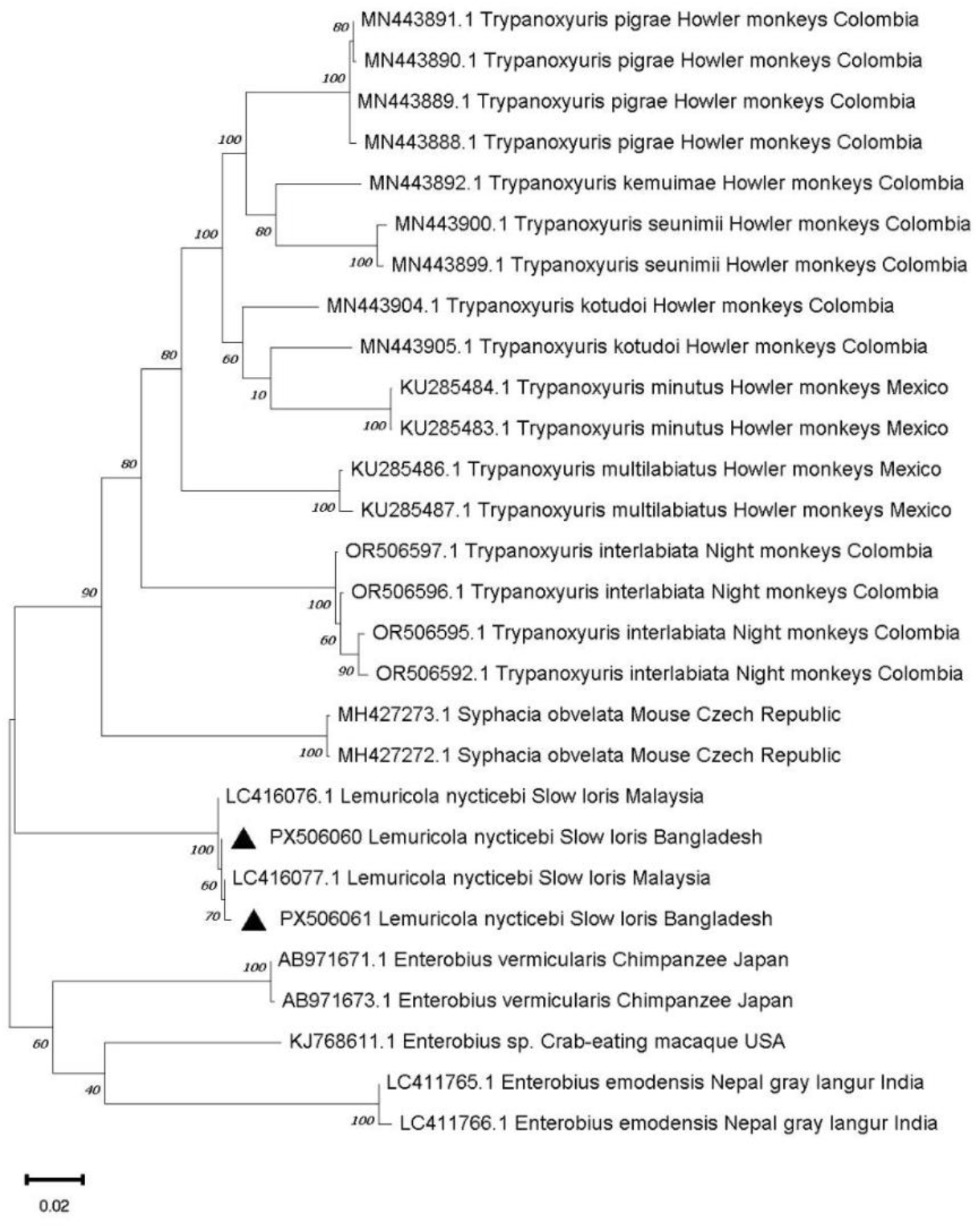
Phylogenetic analysis of query sequences using the maximum-likelihood (ML) algorithm based on *cox1* gene sequences. The numbers at nodes refer to bootstrap values. The sequences generated in this study are highlighted with a black triangle.

## 4. DISCUSSION

The study recovered numerous live adult pinworms from two deceased Bengal slow lorises in northeastern Bangladesh. Morphological traits combined with genetic profiling confirmed the parasites as *L. nycticebi*, and the associated lesions indicate that these infections caused severe pathological damage in both animals. This represents the first confirmed record of *L. nycticebi* infection in the Bengal slow loris and the first evidence of its presence in Bangladesh. Previously, the nearest known occurrence of *L. nycticebi* was documented in Borneo, approximately 3,400 km from our study site [25].

Given this large geographic gap, the route by which *L. nycticebi* entered Bengal slow loris populations remains unclear. The absence of parasitological studies across the species’ range leaves several possibilities open, including natural but undocumented distribution, historical host–parasite associations, or regional transmission dynamics that have not yet been detected. Another plausible explanation is illegal wildlife trafficking, which is widespread in slow lorises across South and Southeast Asia. Over the last two decades, many confiscated or rescued individuals of Bengal slow lorises from different areas were released to Lawachara and Satchari National Parks without adequate health screening [29,31], which could facilitate the movement of infected individuals or parasite lineages across long distances [39]. Hence, without systematic surveillance, the relative contribution of these pathways remains unresolved.

### Pathological consequences

Slow lorises are known to be parasitized by a limited range of gastrointestinal parasites, including hairworms (*Trichostrongylus* spp.), hookworms (*Necator* spp.), spiny-tailed worms (*Pterygodermatites* spp.), threadworms (*Strongyloides* spp.), and tapeworms (*Mathevotaenia* spp.) [22,25,33,40,41,42]. Despite this diversity, *L. nycticebi* remains the only pinworm species reported in lorises to date [22,25]. The prevalence and associated pathology of *L. nycticebi* remain largely unknown in lorises, except for one study reporting a 70% prevalence among 21 wild Javan slow lorises, indicating a high infection rate [22]. Its widespread presence across the peritoneum, intestinal lumen, and peritoneal–diaphragmatic wall in our studied Bengal slow lorises also indicates an unusually intense and invasive helminth infection (Fig. 3). Furthermore, the intestinal hemorrhages and mucosal lesions observed in these animals suggest their substantial migratory and tissue-penetrating capacity and cause acute vascular and epithelial damage. This behaviour is similar to many other invasive helminths reported in wild and captive primates [19,42]. Such helminth infections can lead to chronic nutrient depletion, reduced fitness, and influence survival and reproduction of the host, along with increased susceptibility to secondary infections, including potentially fatal sepsis [19,20,41].

### Conservation Implications

The Bengal slow loris is a globally Endangered species that occurs across northeastern India, Bangladesh, Myanmar, southern China, Laos, Thailand, and Vietnam, but populations are increasingly fragmented and threatened by habitat loss, electrocution, road mortality, hunting, and illegal wildlife trade [23, 27]. The pathology associated with *L. nycticebi* in our specimens suggests that parasitic infection may represent an additional health stressor both in wild and captive populations. The substantial geographic separation between previous *L. nycticebi* records and Bangladesh also raises the possibility of undocumented host movement, including that associated with illegal wildlife trafficking. Given that many Bengal slow lorises are routinely rescued or confiscated from the wildlife trade and forest-edge areas in Bangladesh and subsequently released into the forests [31], our results underscore the need for systematic health assessments, including parasitic screening, before reintroduction.

## 5. CONCLUSION

This study provides the first confirmed record of *L. nycticebi* in the Bengal slow loris and documents associated severe pathology. The new geographic record from Bangladesh highlights an overlooked parasitic risk and the need for broader surveillance. Because many lorises are confiscated and released, routine parasitic screening and health assessments of these animals are essential for informed conservation management.

## Acknowledgement

We gratefully acknowledge the Bangladesh Forest Department for their support and approval to conduct work at the Jankichara Wildlife Rescue and Rehabilitation Center. Our sincere gratitude goes to the Divisional Forest Office, Wildlife Management and Nature Conservation Division, Moulvibazar, for handing over deceased animals for post-mortem at the Jankichara Wildlife Rescue and Rehabilitation Center. We thank the field staff of Creative Conservation Alliance (CCA) for sharing valuable insights and facilitating the work.

